# Attracting Cavities 3.0: Faster and More Versatile Molecular Docking for the SwissDock Webserver

**DOI:** 10.64898/2026.04.21.719847

**Authors:** Ute F. Röhrig, Marine Mathieu-Bugnon, Vincent Zoete

## Abstract

**Motivation:** Molecular docking is a pillar of structure-based drug design and shows advantages in structure prediction of small-molecule ligand–protein complexes over co-folding methods for novel ligands and novel binding pockets. Here, we describe substantial improvements of our physics-based docking algorithm Attracting Cavities, which is widely used through the SwissDock webserver.

**Results:** AC 3.0 includes enhanced sampling features, new functionalities, and technical improvements. These lead to better sampling at lower execution times and higher versatility. Comparison with AutoDock Vina demonstrates better docking results on multiple test sets.

**Availability:** AC 3.0 will be made available free of charge through the SwissDock webserver (www.swissdock.ch).

## Introduction

Molecular docking is a computational approach for predicting the most probable binding mode of a small molecule to a macromolecular target, most commonly a protein, but possibly also a DNA or RNA. Docking algorithms, which predict possible structures for ligand–target complexes and usually also estimate the corresponding binding affinities, constitute the cornerstone of structure-based computer-aided drug design. A docking algorithm generally consists of a sampling algorithm, which generates putative ligand binding modes, and a scoring function, which evaluates and ranks them.

Attracting Cavities (AC)[31] is based on the CHARMM molecular modeling suite [4] and force field (FF).[3, 14] For sampling, the rough energy landscape of the macromolecule is replaced by a smooth attractive energy landscape generated by virtual attracting points representing the macromolecular surface, while for scoring the fast analytical continuum treatment of solvation (FACTS) [11] implicit solvent model is used in combination with the FF energy. In 2023, we described an update of the algorithm and the implementation of new features,[21] such as improvements of the sampling phase leading to better sampling and more robust results, the default use of the current CHARMM36 FF, [3, 14] a shared-memory parallelization of the code, and the possibility to define parts of the receptor as flexible. Additionally, we implemented a covalent docking procedure in AC, mimicking the physical two-step process of non-covalent attraction followed by covalent bond formation, [8] and the possibility of doing on-the-fly hybrid quantum/classical (QM/MM) docking for non-covalent and covalent ligands. [9] Since April 2024, AC 2.0 is deployed on the SwissDock webserver (www.swissdock.ch) [6] and has been used for more than 200,000 docking calculations submitted by users worldwide. Ligand parameters and topologies can be obtained through the SwissParam approach [5] based on the Merck molecular force field (MMFF) [12] or the multipurpose atom-typer for CHARMM (MATCH) [29] approach.

Here, we describe new sampling features in AC, namely (1) the clustering and selection of placement points and (2) the pre-generation of diverse ligand conformations. We also implemented new functionalities, including (1) an active site focus for sampling, (2) stereochemistry checks, (3) pose strain calculations, (4) the option to rescore provided poses with or without relaxation, and (5) settings optimized to dock the heme co-factor. Additionally, technical improvements comprise (1) better CIF/mmCIF file format handling, (2) better compatibility with MATCH parameters, (3) easier covalent docking setup, (4) a check of different cavity prioritization values, the automatic determination of heme-binding potentials, [20] and symmetry-corrected RMSD calculations with spyrmsd [16] for clustering and filtering.

We developed these new features using a part of the PDBbind Core set v2016 [15, 25] taking its known shortcomings into account [21] and evaluated them on the Astex Diverse Set, [13] the Runs N’ Poses (RNP) complex set, [24] and a covalent test set. [8, 9] As we do not modify the scoring function, no extensive benchmarking of the scoring and ranking power of AC is presented. Instead, the conclusion drawn with AC 2.0 remain valid. [21, 8, 9, 23] The popular state-of-the art docking code AutoDock Vina (Vina) [26, 7] was used for comparison with AC. We previously published an in-depth comparison of ligand– protein structure prediction by AC and Vina to AlphaFold 3, [1] using the annotated Runs N’ Poses (RNP) benchmark set. [24] The results demonstrated that Vina and especially AC are advantageous for novel ligands and novel binding pockets when compared to co-folding predictions. [23]

## Methods

### Data Sets

For all datasets, we prepared all structure, topology, and parameter files with the latest versions of SwissParam [5] and CHARMMER, our in-house scripts to prepare CHARMM input files. At variance with earlier work, [21] receptor structures were not minimized, only clashes were removed. This leads to slightly inferior success rates, but minimizes bias and corresponds to the procedure on SwissDock. [6] Structures were visualized and analyzed with UCSF Chimerax. [18]

### PDBbind Core Set

The PDBbind v2016 Core set consists of 57 target clusters with 5 complexes each for a total of 285 ligand–protein complexes. [25] We corrected some issues of the provided data [21] in our own manually curated version. To thoroughly evaluate the AC sampling procedure, we chose a subset of 100 complexes (PDBbind-Sampl) which were previously shown not to be scoring failures and to be sometimes but not always sampling failures.

### Astex Diverse Set

The Astex Diverse set [13] is a set of 85 structures developed for the validation of protein–ligand docking performance with an emphasis on including diverse enzyme classes as well as diverse and drug-like ligands. Here, we used the manually curated structures from our previous studies. [9, 19]

### Runs N’ Poses Set

The Runs N’ Poses (RNP) benchmark set consists of 2,600 protein-ligand systems annotated by their similarity to the training data of different co-folding methods. [24] We previously highlighted imbalances in the RNP set and proposed a higher quality filtered subset of 919 complexes (RNP-F). [23, 22]

### Covalent Set

The CSKDE56 set is a diverse high-quality covalent complex set, including protein–ligand bonds through Cys, Ser, Lys, Glu, and Asp residues formed by eight distinct chemical reactions. [9, 19]. It is part of a larger set of 304 complexes that was used to compare the covalent docking feature of AC with AutoDock and GOLD. [8]

### Docking with AC

All docking calculations were carried out with the CHARMM36 force field [3, 14] and the CHARMM program, [4] version 48b1, removing all solvent molecules during setup, keeping the protein rigid, and using the AC scoring function (total energy with FACTS solvation [11]). Ligand topologies and parameters were generated with the MMFF-based [12] SwissParam approach. [30, 5] For the Astex and PDBbind sets, we also tested MATCH parameters [29] generated with the SwissParam webserver. [5] A randomized ligand conformation was used as input, and the cubic search box center was defined as the center of mass of the ligand in the corresponding experimental structure. The number of saved poses depends on their diversity and was maximally 160 (20 clusters of 8 poses each). All calculations were performed on an Intel Core i9 4.7/3.8 GHz CPU.

By default and unless otherwise stated, all docking calculations were carried out with MMFF-based ligand topologies and parameters, a Morse-like metal binding potential (MMBP) in case of heme binding, [20] a cubic search box with an edge length of 25 Å, an active-site focus without extension of the search box, an attractive threshold value *N*_*Thr*_ of 70, a rotational angle of 90°, 4 random initial conditions (RIC) with defined random seed, 8 placement points (PP), and 4 inital ligand conformations generated with RDKit (version 2025.3.3). The RMSD value to the reference ligand, the SwissParam score, [30] the ligand burial, strain energies, and stereochemistry checks were always calculated. Based on our findings, we call these sampling parameters “standard” parameters (90° rotation, 4 RIC, 8 PP, 4 conformers) and define also “fast” parameters (90°, 2 RIC, 4 PP, 4 conf) and “optimized” parameters (1–20 ligand heavy atoms: standard parameters; 21-35 atoms: 90°, 8 RIC, 4 PP, 4 conf; more than 35 atoms: 90°, 4 RIC, 1 PP, 32 conf).

### Docking with AutoDock Vina

The free and open-source docking program AutoDock Vina, version 1.2.3, [26, 7] was used for comparison to AC. AutoDock Tools (ADT) [17] was used to generate the structure input files for proteins and ligands in pdbqt format starting from the respective CHARMM input files. The search space was defined as for AC, i.e. a cubic box with an edge length of 25 Å an exhaustiveness parameter of 100, and the Vina score were used. For each docking, a maximum of 100 poses was saved (num modes), setting the energy range to a large value. All calculations were performed on a single Intel Core i9 4.7/3.8 GHz CPU.

### Docking Success Criteria

We calculated symmetry-corrected RMSD values between all ligand poses and between the poses and the X-ray structure using the open-source spyrmsd approach. [16] To assess the re-docking performance of different parameters and programs, we analyzed if the experimentally determined (native) pose was (i) within 1.0 Å of the top docking pose; (ii) within 1.5 Å of the top docking pose; (iii) within 2.0 Å of the top docking pose; and (iv) within 2.0 Å of one of all final docking poses. We also calculated and analyzed the median top pose RMSD.

## Results and discussion

### Improvement of Sampling Procedure

#### Previously Implemented Sampling Parameters

In AC 2.0, there were mainly two parameters to determine the sampling exhaustivity, i.e., the ligand rotational angle, a divisor of 360, and the number of random initial conditions (RIC, Fig. 1). Both parameters lead to a better sampling and higher accuracy with increasing CPU time. However, there was a limited flexibility, especially since the minimal number of rotamers per point are 1 (360°), 4 (180°), 24 (90°), 27 (120°), or 108 (60°), with no values in-between. To gain more flexibility and to improve sampling with respect to computational time, we implemented new sampling options in AC 3.0.

**Figure 1.**
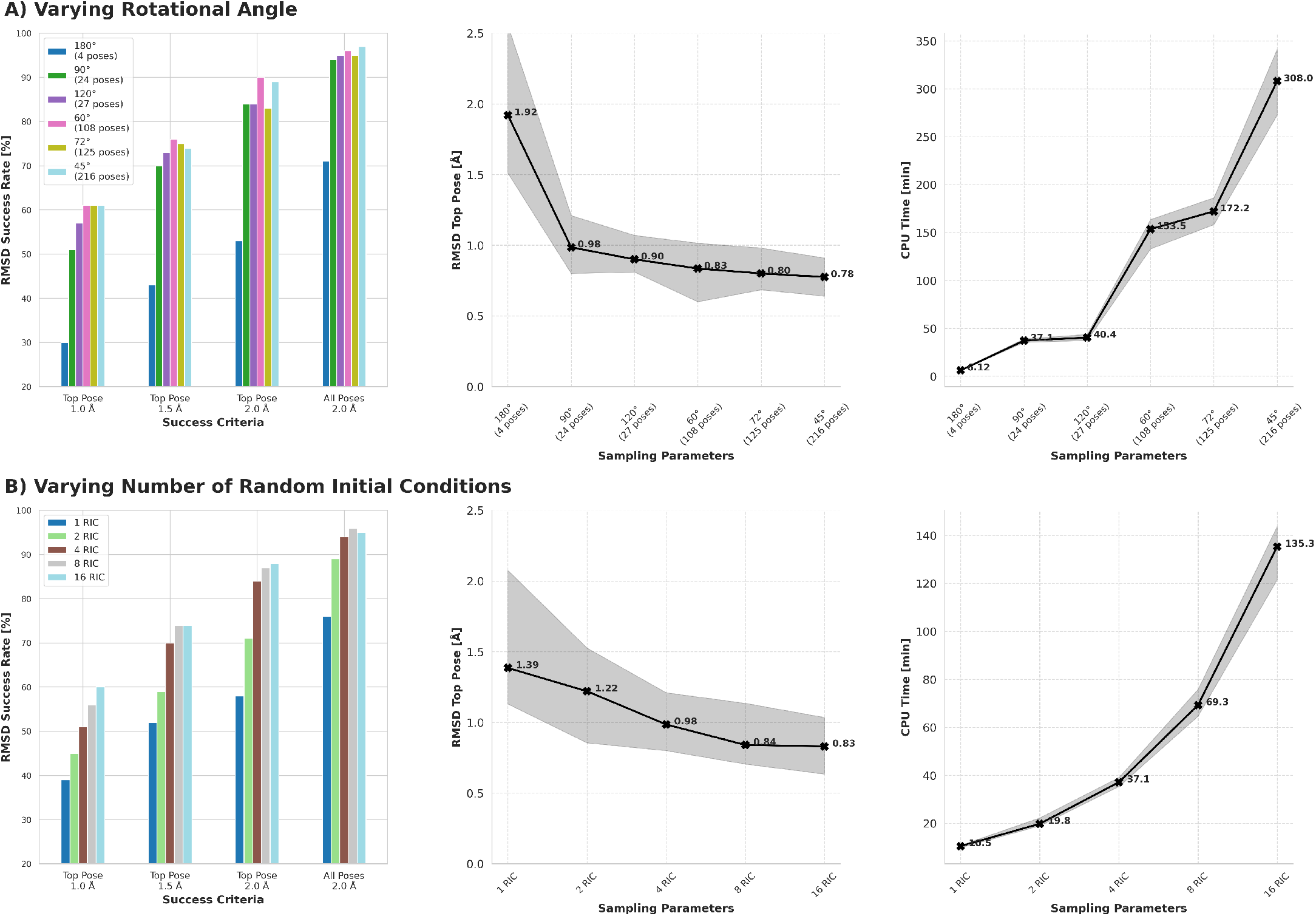
Docking results as a function of sampling parameters implemented in AC 2.0. A) Success rates, top pose RMSD, and CPU time as a function of varying the initial rotational angle of the ligand. B) Success rates, top pose RMSD, and CPU time as a function of varying number of RIC (1–16).

#### Selection of Placement Points

In AC 2.0 we introduced placement points for initial ligand centering, which were defined analogously to the attractive points, but less numerous because removing points close to the protein surface. [21] In AC 3.0, the user can chose the maximal number of placement points to be used for sampling (max_*PP*_). If only one placement point is retained, the geometric center of all determined placement points (tot_*PP*_) is chosen. If only a few placement points are retained, outliers are removed before clustering and retaining cluster centers. If more placement points are retained, clustering is performed without outlier removal. If max_*PP*_ is larger than half of the number of placement points, clustering is skipped and all points are retained.

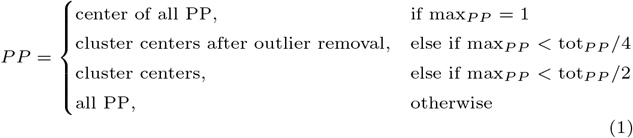

We observed that the success rates significantly increased with an increasing number of placement points (Fig. 2A), but a good sampling was reached with 8 PP. The computational time increased sub-linearly, because of cases with less than the requested number of PP. However, the use of all attractive points as placement points still outperformed all other approaches, but at longer CPU times.

**Figure 2.**
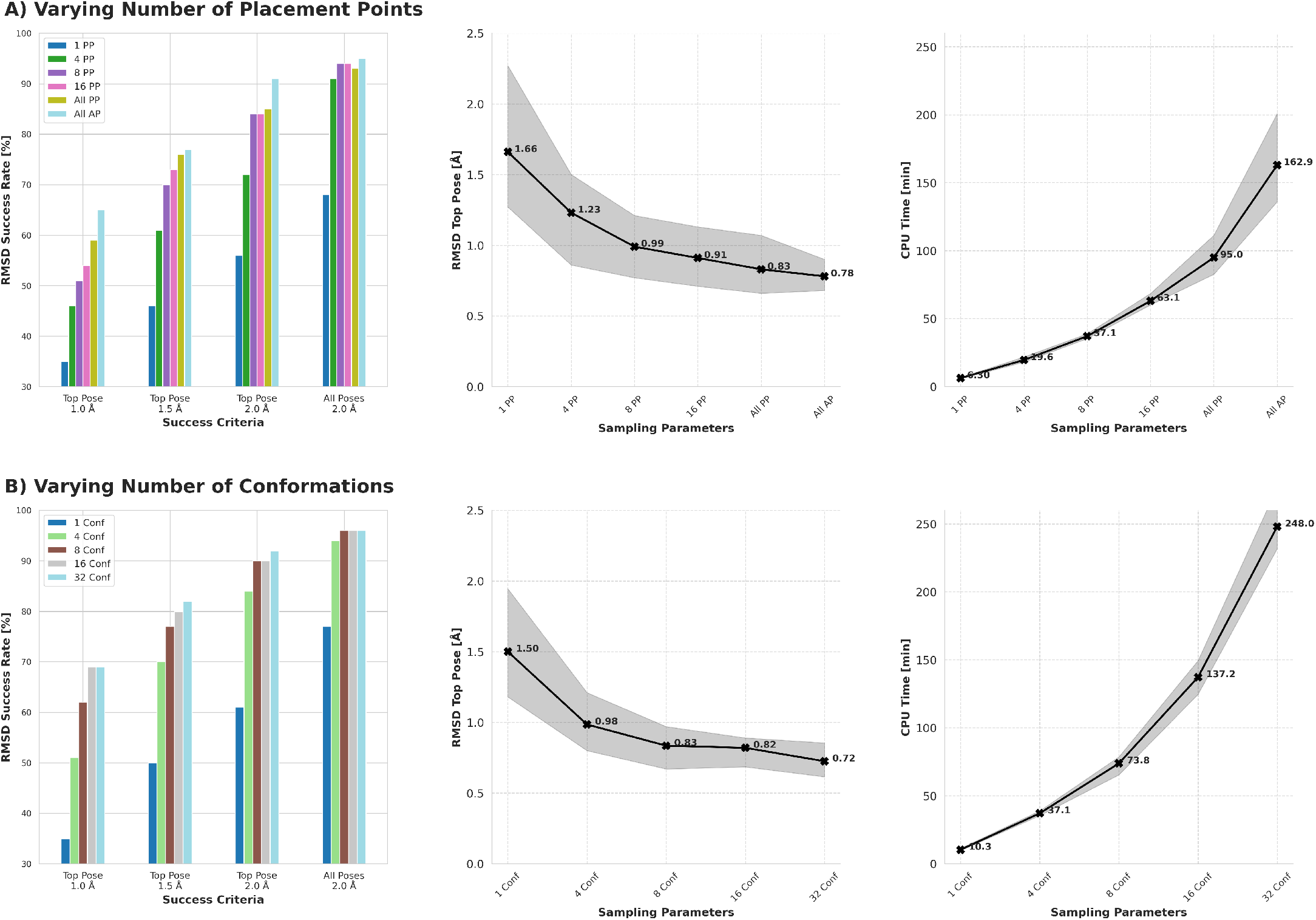
Docking results as a function of new sampling parameters. A) Success rates, top pose RMSD, and CPU time as a function of varying number of placement points. B) Success rates, top pose RMSD, and CPU time as a function of varying number of initial conformations (1–32).

#### Ligand Conformer Generation

In AC 2.0, docking started from one initial ligand conformation, which was stretched to an extended form. In AC 3.0 we implemented the option to generate multiple initial ligand conformations using the *AllChem*.*EmbedMultipleConfs* function of RDKit (version 2025.3.3) with a defined random seed to obtain deterministic results. This option necessitates to provide the ligand input structure in mol2 format, as the pdb format is lacking chemical bond information. The minimal RMSD between accepted conformers was set to 0.1 Å. As anticipated, the success rates on the PDBbind-Sampl set of complexes significantly increased with an increasing number of conformations (Fig. 2B), while the computational time increased.

#### Evaluation of Improved Sampling Procedure

With the described options at hand, in addition to the initial ligand rotational angle and the number of RIC, it is now possible to thoroughly control the number of poses sampled in a docking run with AC 3.0. As the number of poses is the product of the poses generated with each of the 4 sampling options (rotamers, RIC, PP, conformers), care should be taken to not generate an excessive number of poses. We carried out tests to (1) compare efficiency of the new sampling options with the old ones and (2) determine the best balance between sampling parameters. Regarding the first comparison, we found the new options to provide not only more flexibility but also generally better sampling at similar CPU times (Supp. Mat. Fig. S1). To address the second question, we carried out more than 20 dockings on the PDBbind-Sampl set with different sampling parameters, all theoretically yielding a maximum of 3072 poses (Supp. Mat. Fig. S2). We found that docking success was most sensitive to ligand size and that sampling parameters with a single PP and 16 or 32 conformers performed well for large ligands, while a more balanced mix of rotamers, RIC, PP, and conformers performed best on average. Based on these results, we devised a set of “standard” sampling parameters (90° rotation, 4 RIC, 8 PP, 4 conf), a set of “fast” parameters, and and a set of “optimized” parameters, determined during AC execution based on the number of ligand heavy atoms (see Sec. 2.2).

### New Functionalities

#### Active-Site Focus

In previous versions of AC, the full receptor was used during sampling and scoring, which is time-consuming and inefficient especially in case of large receptors. Therefore, we implemented and tested the approach to include only residues having at least one atom within the search space during the pose refinement steps. Optionally, the receptor box size for including residues can be extended with respect to the search space. However, for the scoring step the full receptor is included by default to ensure independence of the score on the chosen box size and position. When re-introducing the full target for scoring, ligand poses clashing with target atoms are removed. We tested this approach on the PDBbind-Sampl set of 100 complexes and found it to be well-suited for accelerating docking with AC without significantly impacting success rates (see Supp. Mat. Fig. S3).

#### Stereochemistry Checks

In addition to improving the sampling, we added new options to AC 3.0, such as the possibility to check and filter ligand poses by their correct stereochemistry. In the past, we assumed that ligand sampling and pose refinement do not modify the stereochemistry of the input conformation of the ligand. This is true as long as covalent ligand bonds are preserved throughout the docking. However, when re-introducing the receptor atoms after the initial ligand sampling in the attractive potential, clashes between ligand and receptor atoms can cause very distorted ligand conformations, leading to a different stereochemistry after relaxation.

In AC 3.0, the bond and atomic stereochemistry of the input conformation is determined by default using RDKit, and each stereoelement of every final pose of a docking run is checked against this reference. If a pose differs in its stereochemistry from the reference, a flag is printed to the output file in DOCK4 format, and the pose can optionally be removed from the results. Both chiral atoms and double bond stereochemistry are taken into account. About 60% of the ligands of each set include at least one stereoelement, mostly a chiral carbon atom (Supp. Mat. Tab. S1) Without surprise, the covalent CSKDE56 set shows the highest percentage of wrong stereochemistry among the top poses (9%) with standard sampling parameters. However, success rates remain very similar when removing these poses (Supp. Mat. Fig. S4).

#### Pose Strain Calculation

Especially for covalent docking, where a covalent bond is forced to form between the ligand and the receptor, it is helpful to assess the ligand strain energy. [10] To this end, we generate a maximum of 100 conformations of the ligand before the docking calculation, using the RDKit conformation generator without setting a minimal RMSD value between conformers. These conformations are minimized with the FACTS solvation model, and for the conformation with the lowest total energy, both its total energy and its bonded energy per atom (*E*_*B*_, Eq. 2) are saved as reference values. If RDKit fails to generate conformations, only the input conformation is minimized and used as reference.

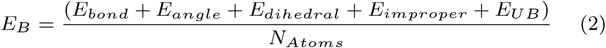

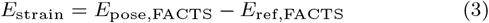

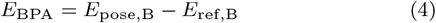

For the final docking poses, the same terms are calculated without minimization, and their differences with respect to the minimized solvated reference conformation are saved. A high strain energy or bonded energy in the top pose suggests caution in the interpretation of the docking results.

#### Rescoring with or without Relaxation

We implemented the possibility to calculate the score for a single or multiple poses, either with or without relaxation. This is useful, for example, if additional terms such as ligand strain, ligand burial, RMSD with respect to a reference pose, or a stereochemistry check should be calculated. Another use case is the assessment of poses generated by a different docking or co-folding algorithm. The input for rescoring can be in PDB, MOL2, or DOCK4 file format, and the output with all calculated terms will be in DOCK4 format.

#### Treatment of Hemoproteins

In AC 3.0, ligand–heme binding potentials [20] can be determined on-the-fly if a heme co-factor is detected inside the search space. We also determined docking parameters optimized for heme co-factor docking to apo-proteins, when only the center of the heme co-factor is known or has been predicted. To accelerate docking of the relatively rigid heme co-factor, one RIC, one PP in the center of the search space and one initial conformer are used, while a rotational angle of 60° allows for sufficient ligand sampling. Application of this approach will be discussed in a forthcoming manuscript.

### Technical Improvements

#### CIF/mmCIF File Format Handling

Recently added PDB entries containing a new ligand are no longer available in PDB file format but only in PDBx/mmCIF format due to the necessary expansion of chemical IDs from 3 to 5 characters. The AC setup scripts are able to handle files in mmCIF format; however, as AC internally relies on pdb format for technical reasons, they will shorten 5-character ligand IDs to unique 4-character IDs. For receptor structures in mmCIF format, our CHARMMER setup scripts uses the mmCIF parser from PDBeCIF [27] and yields valid CHARMM input files even when the receptor includes two or more distinct ligands with similar 5-character ligand IDs. We ran the 49,550 compounds from the Chemical Compound Dictionary (as of 2026-02-25) through the MMFF-like procedure of the SwissParam approach and obtained 47,147 valid topologies and parameters (95%).

#### Compatibility with MATCH Ligand Parameters

The SwissParam server [5] permits ligand parameterization not only based on the MMFF approach, [12] which we use by default due to its robustness and availability for a large variety of chemical functions, but also based on the MATCH approach. [29] In AC 2.0, there were potential atom naming conflicts when using MATCH parameters for the ligand, in case the receptor included a co-factor described with the CHARMM general force field (CGenFF). [28] These issues were resolved in AC 3.0, permitting the use of MATCH parameters. However, for the Astex set, MATCH parameters could only be obtained for 86% of ligands, and for the PDBbind set for 74%. Docking results with standard sampling parameters were slightly inferior with MATCH parameters (Supp. Mat. Fig. S5).

#### Improvements in Covalent Docking Setup

Since version 2.0, AC contains covalent docking options. [8] In the current version, we simplified covalent docking setup, so that during ligand setup it must be specified which atom forms a covalent bond to what type of amino acid side chain and through which chemical reaction. SwissParam then creates all MMFF-based files necessary for covalent docking, compatible with all standard receptor files. In the docking input, the user simply defines the reactive protein atom and the path to the generated ligand parameters. We tested this new procedure on the CSKDE56 set and obtained results in agreement with AC 2.0 (Supp. Mat. Fig. S4). Additionally, stereochemistry checks and ligand strain calculation now facilitate the evaluation of covalent docking predictions.

#### Check of Cavity Prioritization Value

The threshold value for cavity detection (N_*Thr*_) determines which cavities are considered for docking; a N_*Thr*_ of 70 detects mainly deep binding clefts, while a value of 60 places attractive points also in medium and a value of 50 also in shallow receptor cavities. [31] The optimal value depends on the receptor and on the chosen search space size, and can be checked by printing and visualizing the attractive points. With the large sampling box of (25 Å)^3^ used in this work to reduce the re-docking bias, generally a N_*Thr*_ value of 70 worked best (Supp. Mat. Fig. S6).

As a heuristic, we observed that the number of attractive points inside the receptor active site should not be smaller than the number of heavy atoms in the ligand to allow for a good sampling. In AC 3.0, during initialization the attractive points for all three N_*Thr*_ values are determined, and their number is printed. If the number of points at the chosen N_*Thr*_ value is smaller than the number of ligand atoms, a warning is printed, suggesting to visually inspect the attractive points and to adapt the box size and N_*Thr*_ value if necessary.

#### Symmetry-Corrected RMSD Calculations

AC 3.0 relies on the open-source spyrmsd python module [16] to calculate symmetry-aware RMSD values between all ligand poses for clustering and between the poses and a reference structure. This approach yields the same results as the DockRMSD approach [2] used in AC 2.0 [21] and has the advantage to be easily integrated into the docking algorithm for pose clustering, similarity filtering, and RMSD calculation to a reference ligand structure.

### Re-Docking Comparison of AC and AutoDock Vina

We compared the performance of AC 3.0 to the one of AutoDock Vina on three diverse docking test sets. The results for the PDBbind set (Fig. 3), the Astex set (Supp. Mat. Fig. S7) and the RNP set (Supp. Mat. Fig. S8) demonstrate that AC 3.0 consistently outperforms AutoDock Vina at similar CPU times and shows even higher success rates with better sampling at longer CPU times.

**Figure 3.**
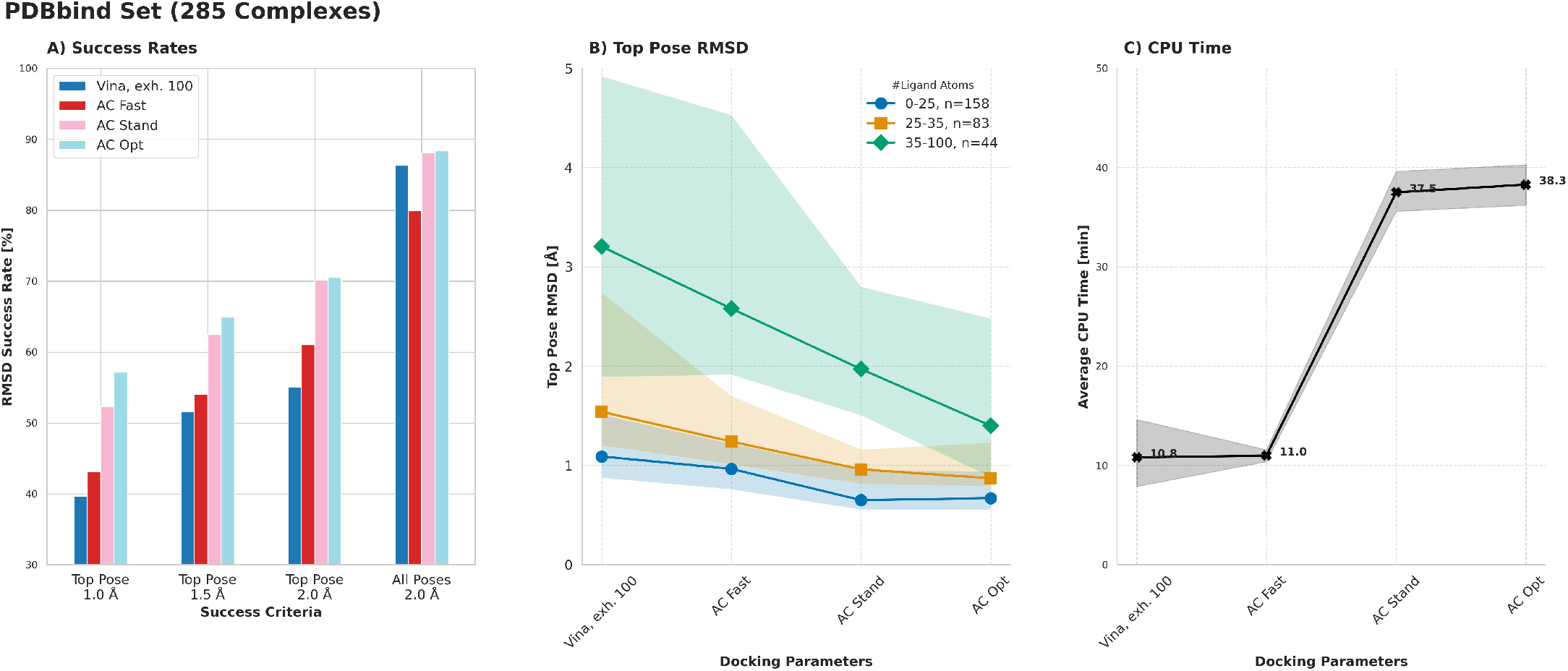
Comparison on docking results of AC 3.0 with AutoDock Vina on the PDBbind Core v2016 set.

## Summary and Conclusions

In summary, version 3.0 of the AC docking algorithm shows significant improvements over version 2.0, including enhanced sampling features, novel functionalities, and technical improvements. None of these developments changes its physics-based scoring function based on the CHARMM force field and the FACTS implicit solvent model, which has proven its value in many applications. However, improved execution speed and novel functionalities broaden the applicability of AC and allow to extend its use on the SwissDock webserver (www.swissdock.ch). [6]

## Data availability

Docking input files, results, and annotations of the present work will be made publicly available on Zenodo upon manuscript acceptance. Access to AC 3.0 will be provided through he login-free SwissDock web server (www.swissdock.ch).

## Author contributions statement

U.F.R. implemented the code, carried out the calculations, analyzed the results, and wrote the manuscript. M.M.B. contributed to the implementation and analysis. V.Z. contributed to the conceptualization and analysis. All authors reviewed the manuscript.

