## Supplementary data for "Attracting Cavities 3.0: Faster and More Versatile Molecular Docking for the SwissDock Webserver"

### Supporting Tables

| Set | Stereo [%] | Chiral carbon [%] | Double Bond [%] | Top Wrong [%] | Max Wrong [%] |
|---|---|---|---|---|---|
| **Astex** | 65 | 56 | 14 | 0 | 13 (1hww) |
| **PDBbind** | 59 | 55 | 9 | 1 | 27 (3g0w) |
| **RNP** | 62 | 59 | 6 | 3 | 87 (7rbw) |
| **CSKDE56** | 63 | 55 | 18 | 9 | 85 (4wsk) |

**Table S1.** Analysis of stereochemistry across data sets. Stereo: ligands presenting at least one stereo element. Chiral carbon: ligands presenting at least one chiral carbon atom. Double bond: ligands presenting at least one stereogenic double bond. The last two columns relate to docking results with standard parameters (90° rotation, 4 RIC, 8 PP, 4 initial conformers). Top wrong: top poses with wrong stereochemistry. Max wrong: PDB ID and percentage of poses with wrong stereochemistry of complex with maximal percentage of poses with wrong stereochemistry. All values are given as percentages.

### Supporting Figures

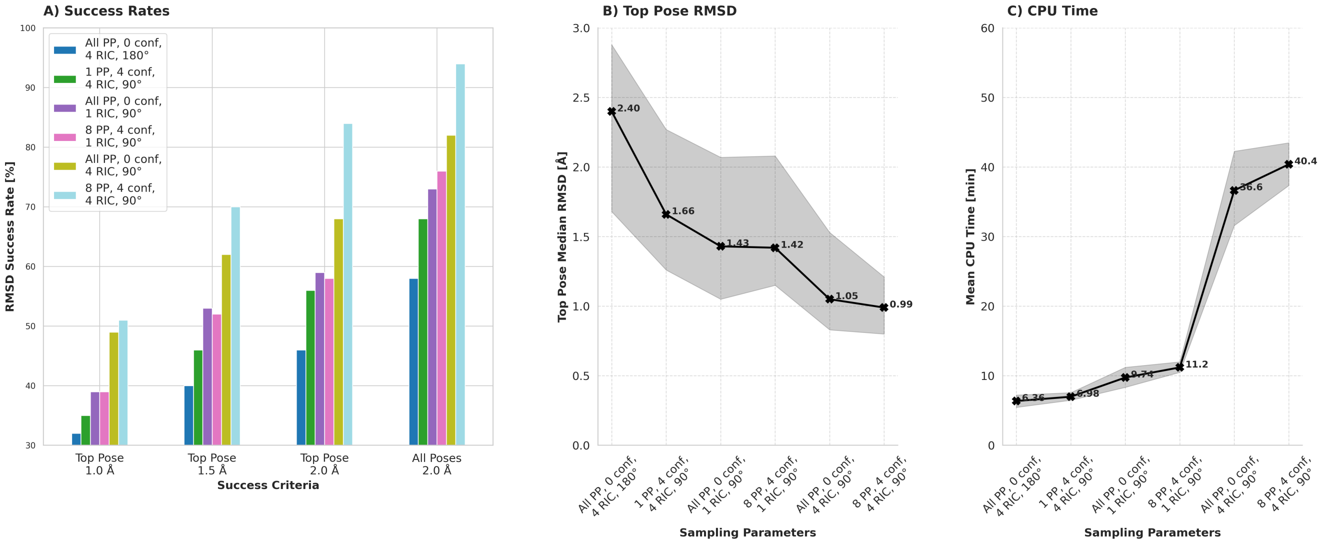

**Figure S1.** AC Docking results on PDBbind-Sampl set of 100 complexes with only old and a combination of old and new sampling options. Dockings with similar mean CPU timings are shown for the two settings. In general, at similar CPU timings better sampling is obtained by integrating the new options.

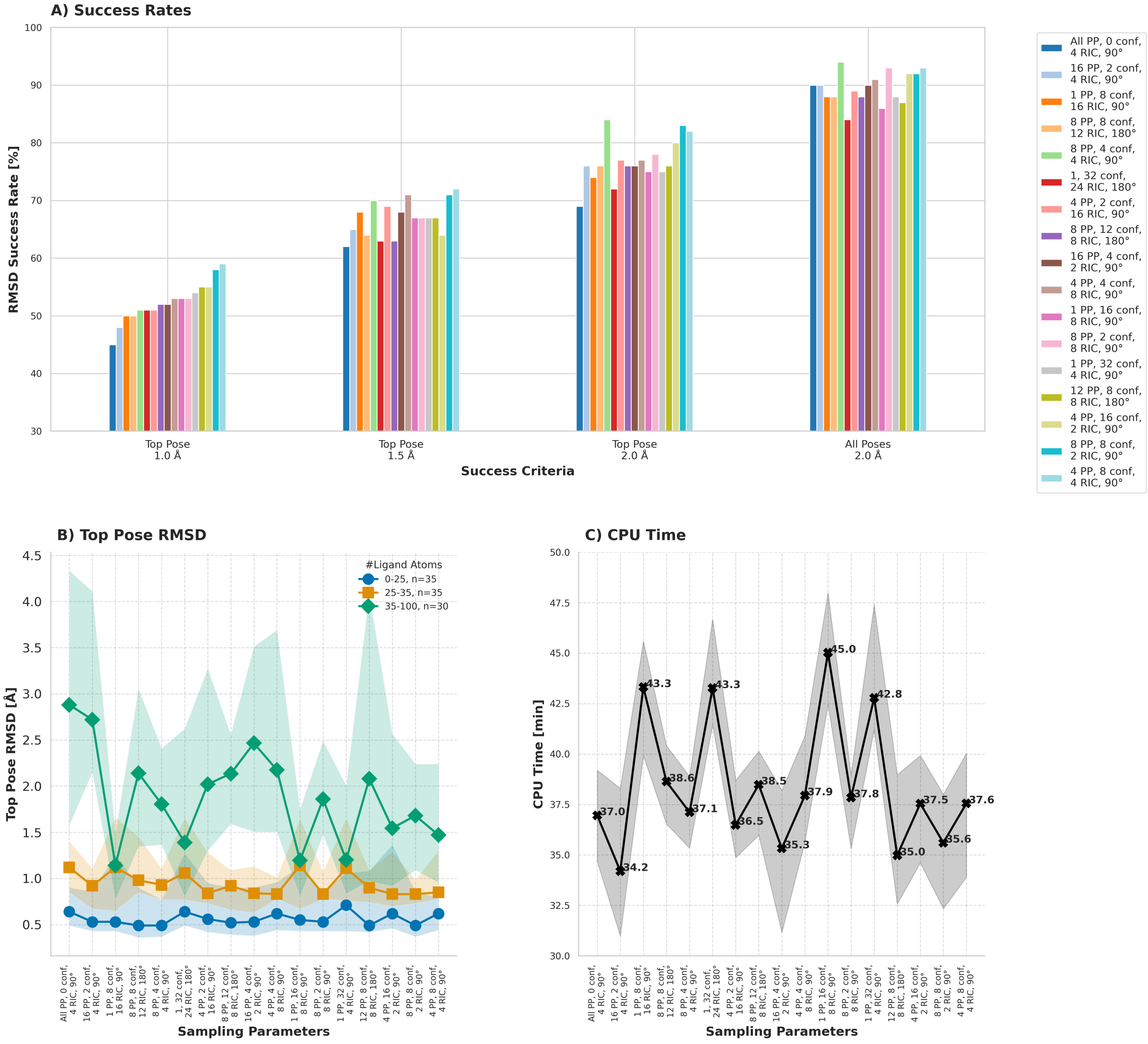

**Figure S2.** AC Docking results on PDBbind-Sampl set of 100 complexes with options theoretically yielding 3072 starting poses. A) Success rates at different RMSD cutoffs. B) Top pose RMSD, displayed by ligand size. C) CPU timings. The biggest differences in sampling were observed for large ligands with more than 35 heavy atoms.

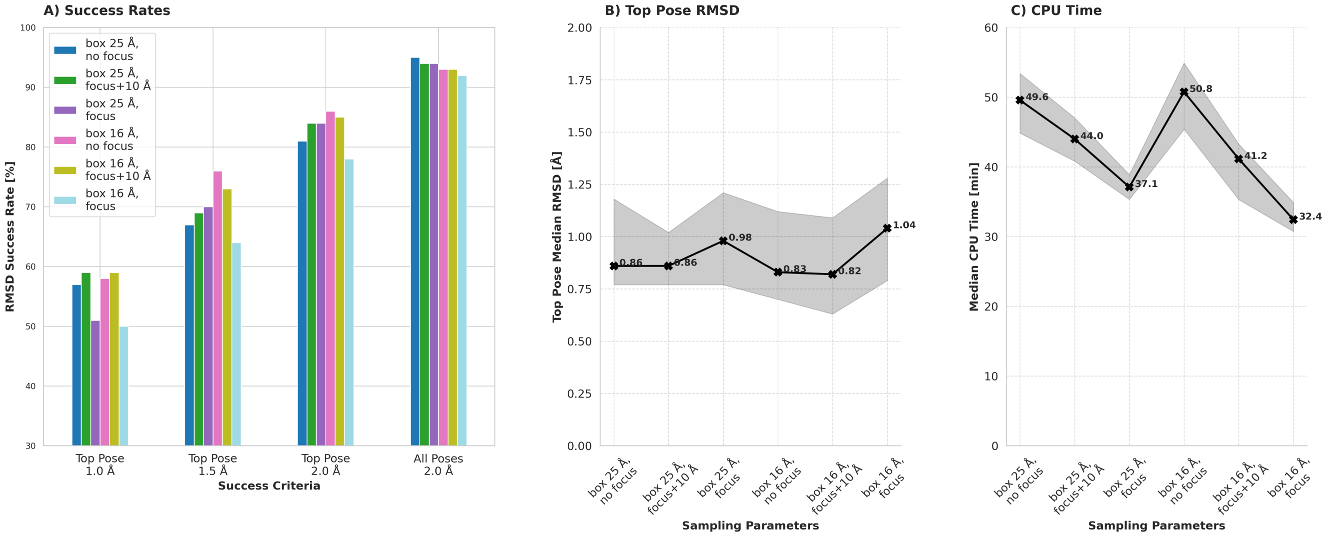

**Figure S3.** Docking results on PDBbind-Sampl set of 100 complexes with and without active-site focus. Two different search space sizes, (25 Å)$^3$ and (16 Å)$^3$ were used. For each size, 3 calculations were carried out, one without focus, one with focus on the search space + 10 Å extension, and one with focus on the box without extension. A focus on the search space reduced CPU times by almost 30% without extension and about half that amount with extension. For the larger box, the success rate at 2.0 Å RMSD and top pose RMSD values remained quite stable, while for the smaller box, a drop in successful docking predictions can be observed.

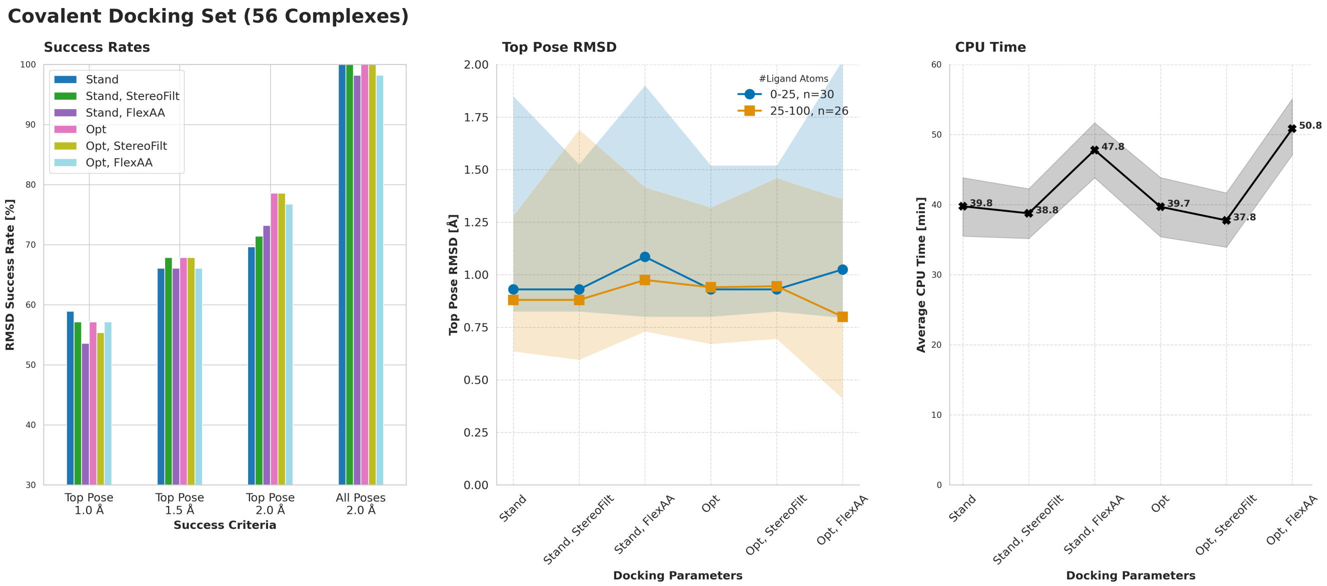

**Figure S4.** Covalent docking results with standard and optimized sampling parameters. Additionally, results are shown for docking when filtering our poses with wrong stereochemistry ("StereoFilt") and when leaving the reactive protein side chain flexible ("FlexAA"). Stereochemistry filtering does not show a significant impact on the results, while a flexible side chain leads to slightly worse results. However, this option may be very beneficial in cross-docking settings.

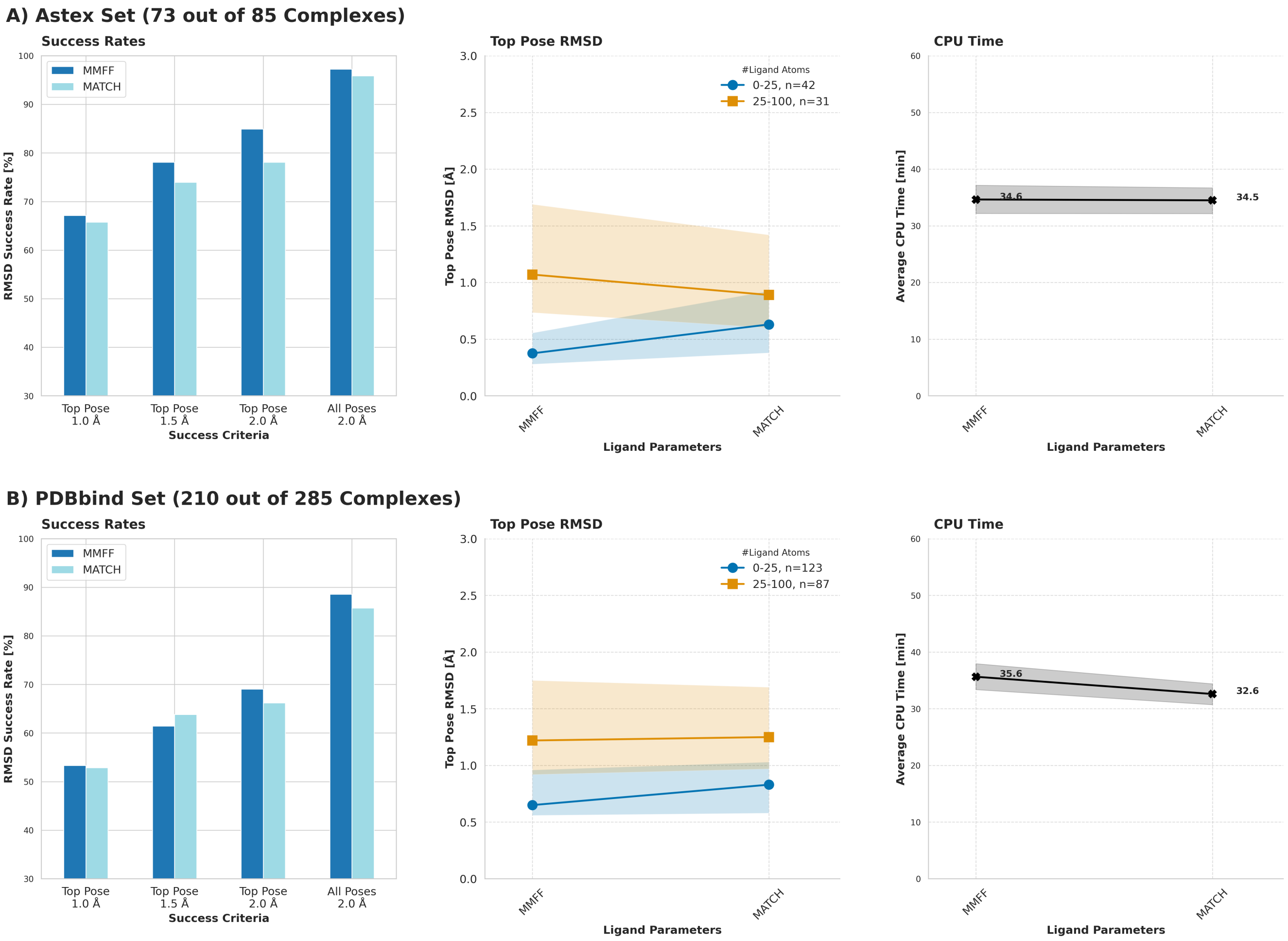

**Figure S5.** Comparison of docking results with standard sampling parameters (90° rotation, 4 RIC, 8 PP, 4 initial conformers) and with ligand parameters generated by MMFF-like[1] and by MATCH[2] approaches. A) Results on subset of Astex set, for which MATCH parameters were obtained. B) Results on subset of PDBbind set, for which MATCH parameters were obtained.

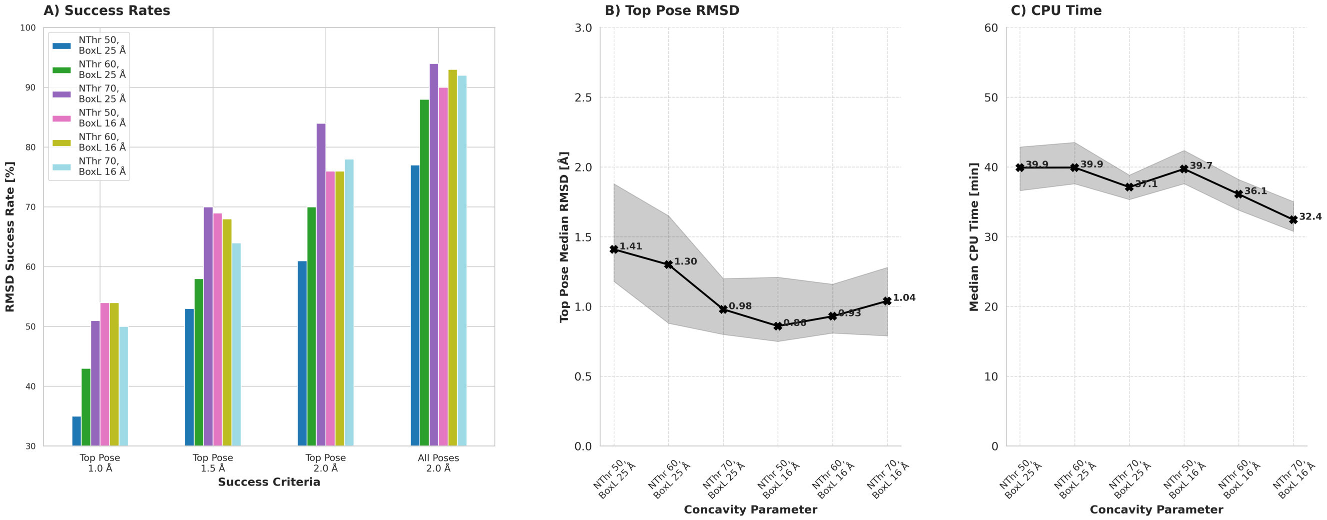

**Figure S6.** AC docking results on PDBbind-Sampl set of 100 complexes with different concavity threshold values. The results demonstrate that the optimal value depends on the search box size.

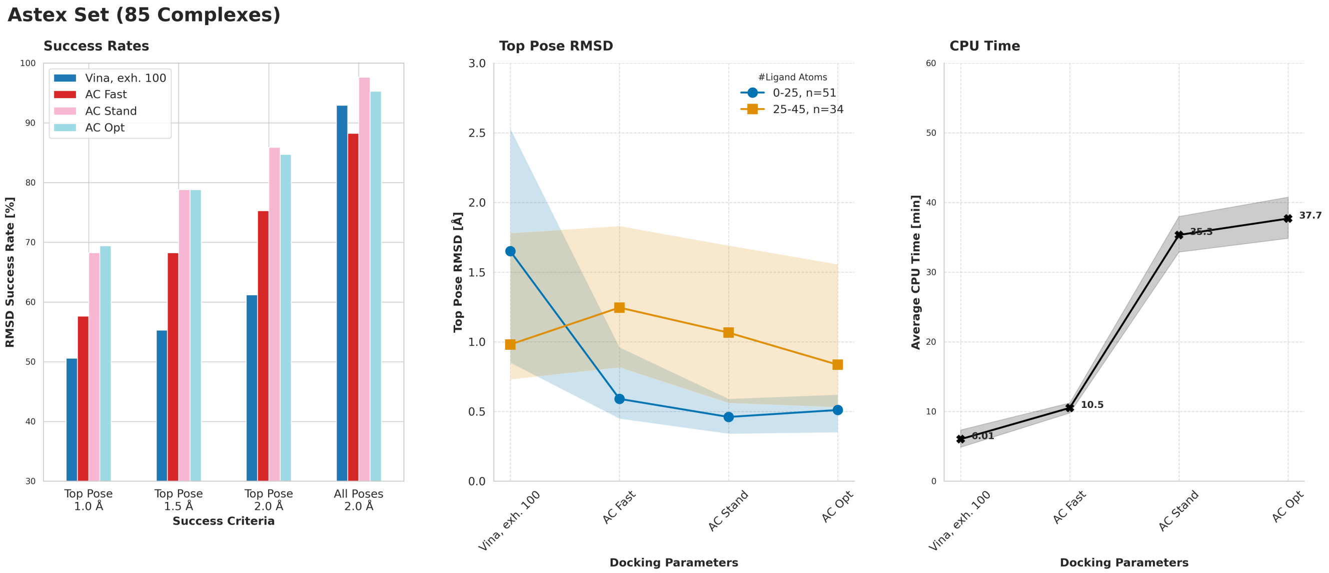

**Figure S7.** Comparison of docking results of AC 3.0 with AutoDock Vina on the Astex Diverse set.[3] A) Success rates. B) Top pose RMSD, displayed by number of ligand atoms. C) Average CPU times.

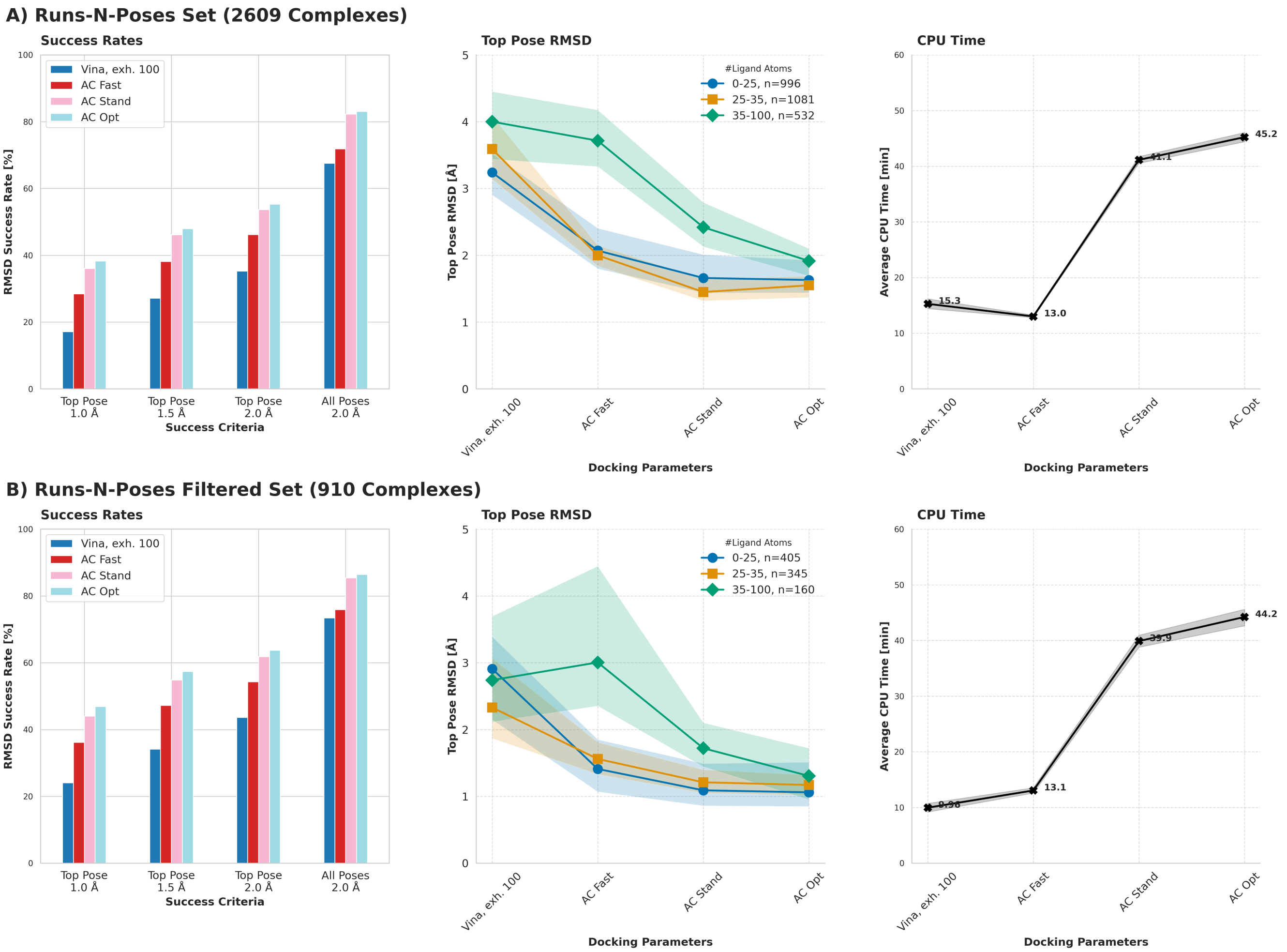

**Figure S8.** Comparison of docking results of AC 3.0 with AutoDock Vina A) on the RNP set[4] and B) on its version filtered for quality and diversity (RNP-F)[5].
